# Rapid development of *E. coli* resistance to *Bdellovibrio* predation during short-term co-culture

**DOI:** 10.64898/2026.11.23.768573

**Authors:** Laura E. Williams, Isabelle Grillo, Katie Mulvey, Caroline E. Zielinski

## Abstract

When *E. coli* ML35 is co-cultured with *Bdellovibrio* sp. NC01 over 72 hours, prey populations initially experience a 3-log reduction, but then partially recover to ∼20% of starting density. Population recovery is correlated with emergence of irregular colonies, which are distinct from typical circular ML35 morphotype. ML35 populations do not recover when co-cultured with *Bdellovibrio bacteriovorus* HD100, and irregular colonies are not observed in HD100 co-cultures or in prey-only controls. Whole-genome sequencing of clonal lines of irregular and circular morphotypes from three independent replicate assays confirmed they are derived from ML35. ML35-irregular is resistant to killing by *Bdellovibrio* sp. NC01 but susceptible to killing by HD100, indicating strain-specific predation resistance. Analysis of sequencing data showed that all three ML35-irregular lines and all three ML35-circular lines carry a mutation in *rfbG*, annotated as dTDP-rhamnosyl transferase involved in O-antigen synthesis. Two ML35-irregular lines have distinct IS element insertions in *rfbG*; the third has a substitution introducing a premature stop codon near the C-terminus. Two ML35-circular lines have the same missense mutation in *rfbG*, whereas the third has a nonstop mutation. Overall, these data suggest that *rfbG* may be under selective pressure during NC01 co-culture and that O-antigen may play a role in predation resistance. Further work, such as phenotypic testing of ML35-circular lines and reverse genetics experiments, are needed to clarify how *rfbG* affects prey susceptibility to *Bdellovibrio* predation.

## INTRODUCTION

As obligate predators of Gram-negative bacteria, including drug-resistant pathogens, *Bdellovibrio* have attracted interest as potential biocontrol agents [1]. Defining how prey respond to *Bdellovibrio* predation, especially during short-term exposure, may inform development of predatory bacteria for clinical and other applications. Susceptibility to predation depends in part on intrinsic features of the prey bacteria. For example, production of cyanide [2] and indole [3] can inhibit predation, whereas outer surface structures such as curli fibers [4] and S-layers [5] can reduce susceptibility to predation by interfering with *Bdellovibrio* attachment. In addition, researchers recently showed that engineered variation in LPS architecture can significantly affect susceptibility to *Bdellovibrio* predation, with some effects differing among *Bdellovibrio* strains [6]. These studies underscore how prey range is shaped by multiple traits and depends on the specific combination of predator and prey strains.

In addition to intrinsic features, changes in prey susceptibility to predation can emerge during co-culture. Early work by Varon described the appearance of predation-resistant colonies when predatory bacteria were introduced into a continuous culture of prey bacteria [7]. Shemesh and Jurkevitch reported recovery of prey populations during co-culture with *Bdellovibrio* and showed that surviving prey were less susceptible to predation [8]. This resistance phenotype was not specific to the predatory strain from the co-culture; however, it was transient and reverted rapidly upon removal of predation pressure, suggesting a physiological response rather than a genetic change. More recently, Huwiler’s research group reported emergence of predation resistance in *E. coli* K-12 following repeated cycles of exposure to *Bdellovibrio* and recovery, with mutations identified in genes associated with the prey cell envelope [9].

Despite these advances, relatively little is known about whether predation resistance that emerges during short-term predator-prey interactions can be predator-specific and what changes are associated with its development. We previously reported that *E. coli* ML35 populations initially decline but then partially recover during 72-h co-culture with *Bdellovibrio* sp. NC01 [10]. Here, we show that prey population recovery is correlated with rapid emergence of a distinct colony morphotype that is resistant to killing by NC01 but remains susceptible to killing by *Bdellovibrio bacteriovorus* HD100. Whole-genome sequencing of both colony morphotypes isolated from NC01 co-cultures identified mutations in *rfbG*, a gene involved in O-antigen synthesis. These findings highlight cell envelope structures as potential contributors to rapid development of strain-specific predation resistance.

## MATERIALS AND METHODS

### Predation efficiency assays

To validate and extend observations originally reported in [10], we repeated the predation efficiency assay described in that study in five independent biological replicates and distinguished colony morphotypes by eye on cfu/ml plates at each time point. During three replicate assays, we used 48-h plates of the ML35+NC01 co-culture to establish independent lines of ML35-irregular and ML35-circular colony morphotypes. To do this, we picked single colonies of each morphotype and streaked to singles on TSA. From these plates, we picked and streaked again, then picked a single colony and inoculated TSB for overnight culture, which we combined with 50% glycerol and stored at -80°C. Colony morphotype remained consistent on all streak plates.

To test *Bdellovibrio* predation efficiency against the ML35-irregular colony morphotype, we first revived NC01 and HD100 by combining a small amount of freezer stock, 20 ml HM buffer, and 1.5 ml *E. coli* ML35 overnight culture and incubating at 28°C and 200 rpm for 72 h. To cultivate the ML35-irregular colony morphotype, we streaked freezer stocks to single colonies on TSA plates and incubated at room temperature for 2 days. We then picked a single colony and inoculated overnight cultures in 15 ml TSB. Following the predation efficiency assay procedure described in [10], we tested NC01 and HD100 against the three independent ML35-irregular lines. We obtained predator concentrations at the start and end of the assay by plaque plating on *E. coli* ML35 prey lawn.

### Genome sequencing and analysis

To obtain genomic DNA for sequencing, we inoculated 3 ml TSB with a small amount of freezer stock from the ML35-irregular and ML35-circular lines established during predation efficiency assay replicates. We incubated cultures overnight at 30°C, then harvested genomic DNA using the Wizard Genomic DNA purification kit (Promega) following the manufacturer’s instructions. At the final step, we rehydrated genomic DNA in nuclease-free water.

We quantified genomic DNA concentration using a Qubit, then diluted with nuclease-free water as needed to yield 35 ng/µl. We shipped dilutions on ice to the Rhode Island Genomics and Sequencing Center (RIGSC) for library preparation and sequencing. After shearing genomic DNA using a Covaris S-220 focused ultrasonicator, RIGSC prepared libraries using the PrepX DNA library kit, visualized the libraries on high-sensitivity BioAnalyzer chips, and quantified them using KAPA Illumina quantification kits. RIGSC sequenced libraries on an Illumina MiSeq instrument using 500-cycle v2 chemistry to generate 2 × 250-bp paired-end reads. Sequencing datasets are available at NCBI Sequence Read Archive under BioProject PRJNA1564189.

We used *breseq* 0.35.5 with default parameters to analyze read datasets by comparing them against the *E. coli* ML35 genome (NZ_CP025747) that we previously sequenced [11].

## RESULTS

### ML35 population recovery during co-culture with NC01 is correlated with emergence of irregular colonies

When we co-cultured *E. coli* ML35 with *Bdellovibrio* sp. NC01, prey populations initially declined by 3.3 ± 0.4 log_10_ cfu/ml within 24 hours, then increased by 2.6 ± 0.5 log_10_ cfu/ml over the next 48 hours, reaching roughly 20% of the starting density at 72 hours (Figure 1). By contrast, when we co-cultured *E. coli* ML35 with *Bdellovibrio bacteriovorus* HD100, prey populations declined by 4.5 ± 0.6 log_10_ cfu/ml within 24 hours and remained at this reduced density at 48 and 72 hours.

**Figure 1.**
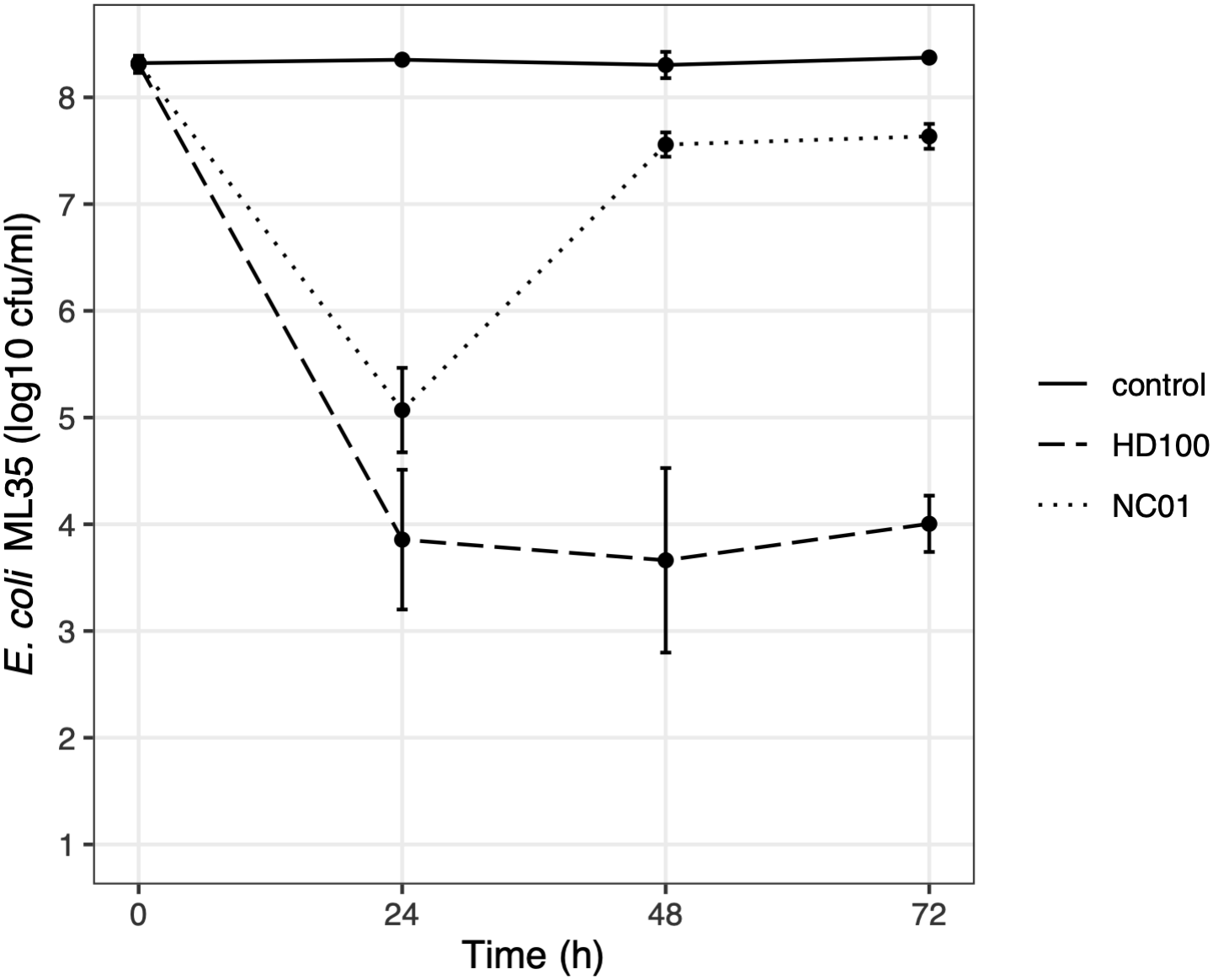
*E. coli* ML35 population recovery during short-term co-culture with *Bdellovibrio* sp. NC01. We incubated *E. coli* ML35 alone (control), with *Bdellovibrio* sp. NC01, or with *Bdellovibrio bacteriovorus* HD100. We counted total colonies to determine prey concentration and plotted the mean and standard deviation of log-transformed counts. All data points represent five independent biological replicates, except the 72-h time point for the control and the 48-h time point for the HD100 co-culture, each of which represent four replicates.

*E. coli* ML35 population recovery during co-culture with *Bdellovibrio* NC01 is correlated with emergence of a distinct colony morphotype. Whereas ML35 typically grows on TSA as circular colonies with entire edges (Figure 2a), we noted the appearance of irregular colonies with undulate edges during co-culture with NC01 (Figure 2b). Across five replicate assays, we always observed irregular colonies at 24 hours but in proportions ranging from 22-75% of total colonies (Figure 2c). At 48 and 72 hours, the irregular morphotype was the predominant colony morphotype. We did not observe the irregular morphotype in prey-only controls or co-cultures with HD100. We established clonal lines of irregular and circular morphotypes from 48-h plates of three independent replicate assays, and whole-genome sequencing confirmed that each line is derived from *E. coli* ML35. We will refer to the irregular morphotype as ML35-irregular and the circular morphotype as ML35-circular.

**Figure 2.**
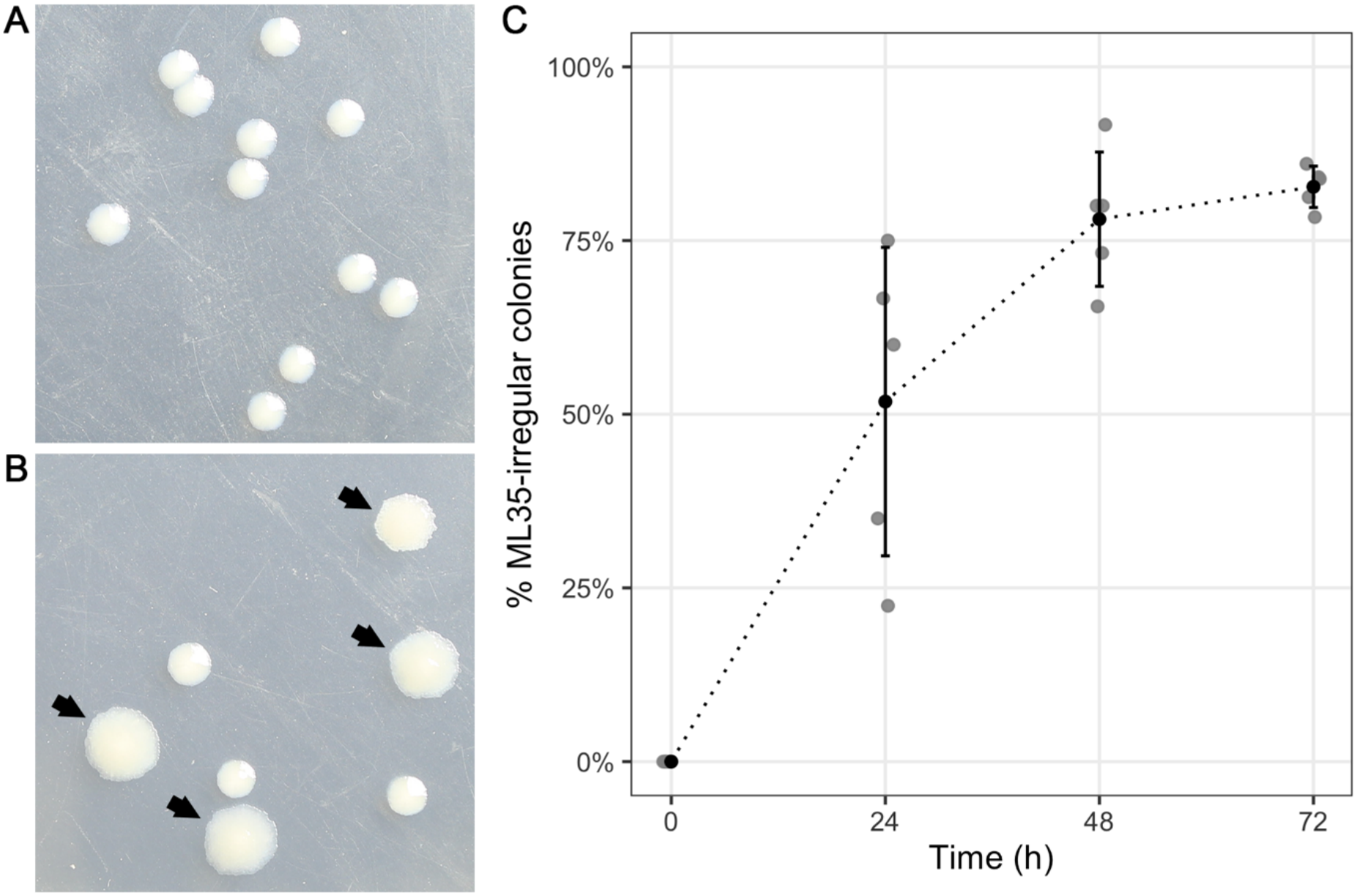
Emergence of ML35-irregular morphotype during short-term co-culture with NC01. (a) Representative ML35 colonies recovered from co-culture with *B. bacteriovorus* HD100 at 48 hours. Colonies are circular with entire edges, which is the characteristic ML35 morphotype (ML35-circular). (b) Representative ML35 colonies recovered from co-culture with *Bdellovibrio* sp. NC01 at 48 hours. Arrowheads indicate irregular colonies with undulate edges, which we refer to as the ML35-irregular morphotype. (c) Proportion of ML35-irregular colonies observed at each time point during short-term co-culture with NC01. Data are from the same five independent biological replicates shown in Figure 1. Grey points represent individual replicates, and black points with error bars show the mean and standard deviation.

### ML35-irregular is resistant to predation by NC01 but not HD100

We co-cultured the ML35-irregular lines with *Bdellovibrio* sp. NC01 and *Bdellovibrio bacteriovorus* HD100 (Figure 3). NC01 failed to reduce ML35-irregular populations over 72 hours, demonstrating that the irregular colony morphotype is resistant to predation by NC01. However, HD100 was able to kill ML35-irregular, further demonstrating that the resistance phenotype is predator-specific. As shown by pfu/ml data (Figure 4), the NC01 live predator population decreased over 72 hours when co-cultured with ML35-irregular, whereas the HD100 live predator population increased.

**Figure 3.**
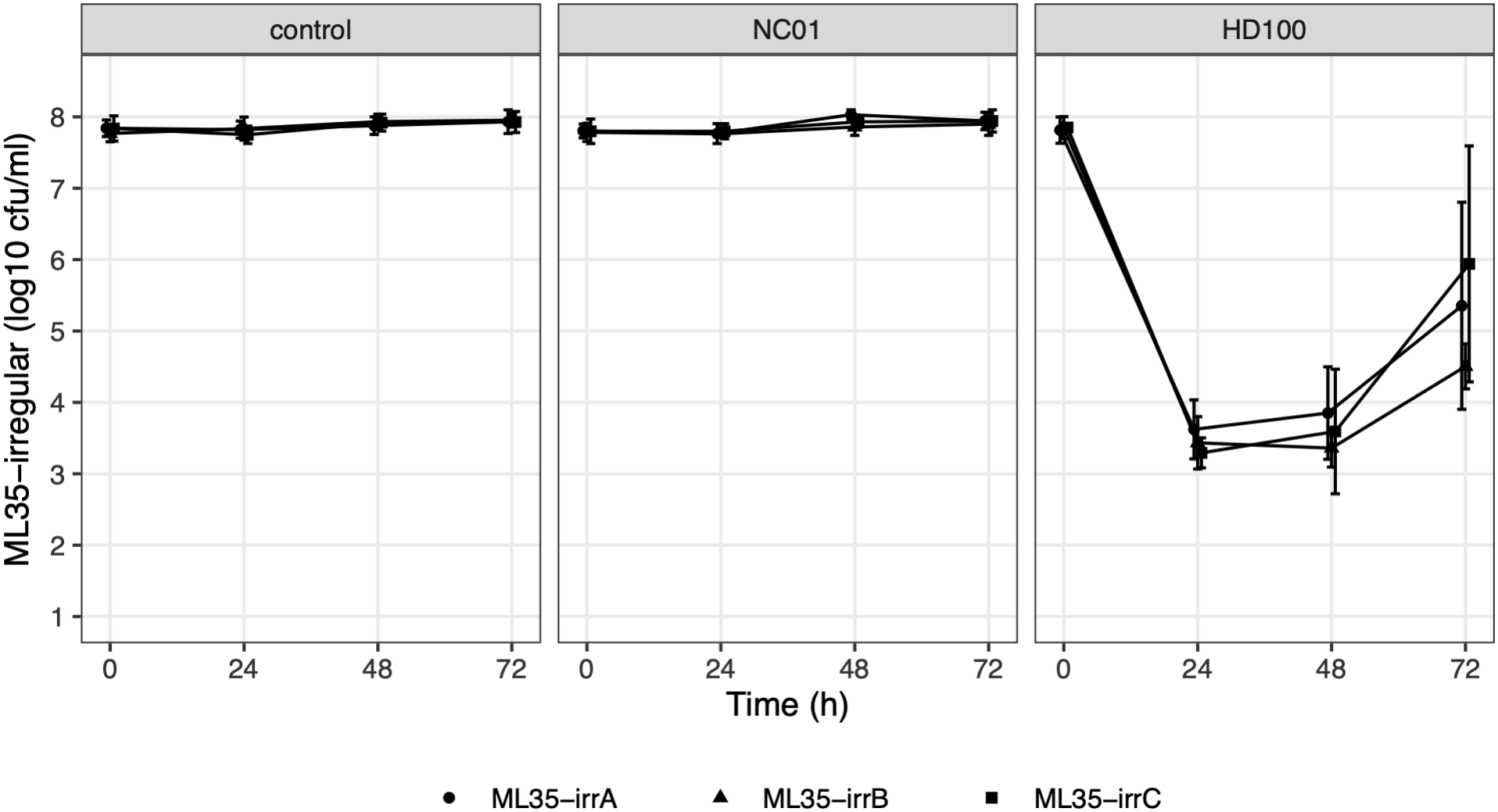
Predation efficiency assays with ML35-irregular as prey. We performed predation efficiency assays using three ML35-irregular lines established from independent NC01 co-cultures (ML35-irrA, ML35-irrB, ML35-irrC) as prey. Panels show prey population dynamics for each ML35-irregular line alone (control), with *Bdellovibrio* sp. NC01, or with *B. bacteriovorus* HD100. We counted total colonies to determine prey concentration and plotted the mean and standard deviation of log-transformed counts. Datapoints for ML35-irrA, ML35-irrB, and ML35-irrC are represented as circles, triangles, and squares, respectively. Each datapoint represents five independent biological replicates, except the 72-h time point for the HD100 co-culture, which represents four biological replicates for each ML35-irregular line.

**Figure 4.**
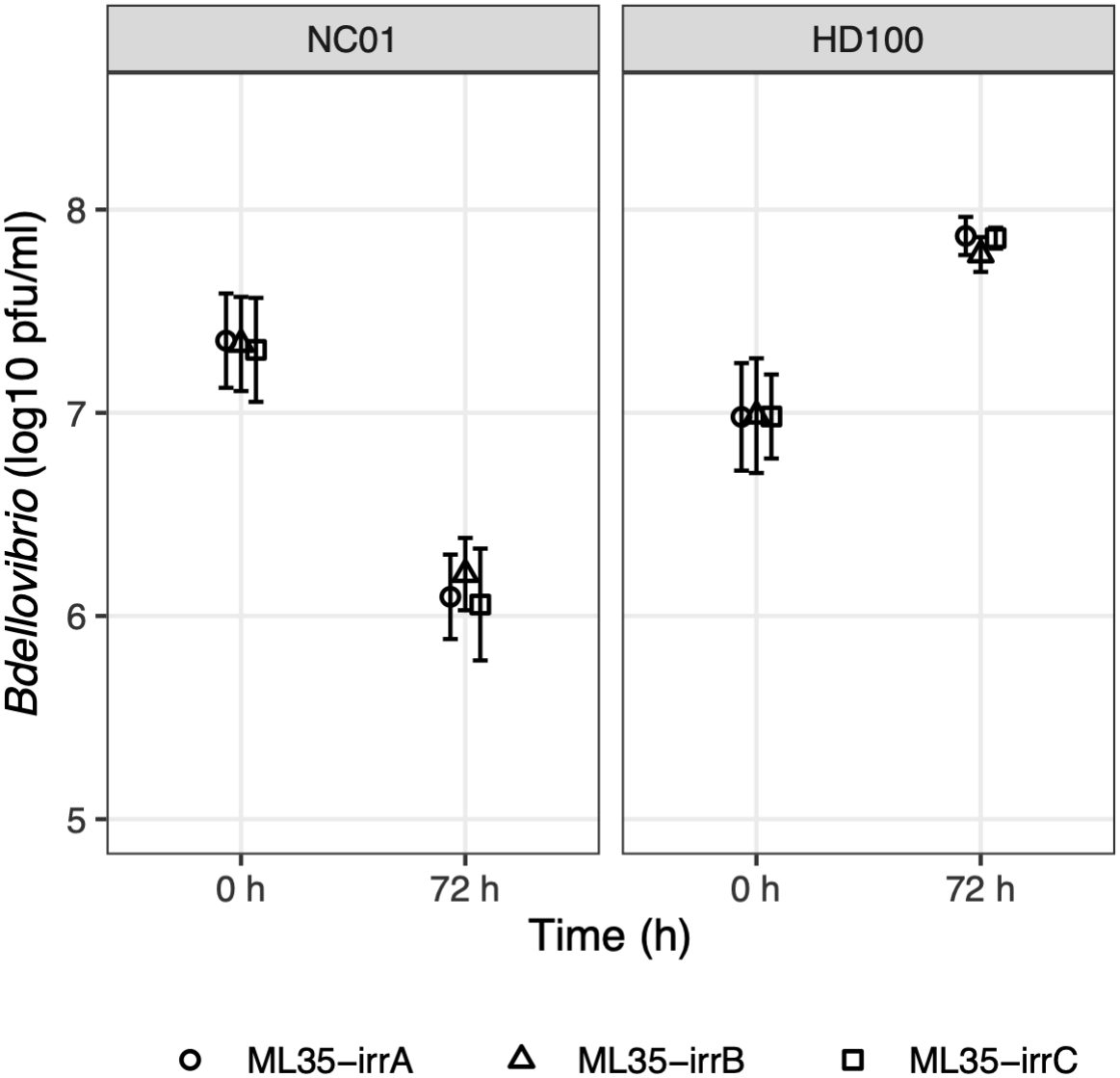
*Bdellovibrio* populations during co-culture with ML35-irregular. We counted plaque-forming units at time zero and 72 hours during the predation efficiency assays shown in Figure 3, with the ML35-irregular lines used as prey in these assays indicated by circles (ML35-irrA), triangles (ML35-irrB), and squares (ML35-irrC). Datapoints show the mean and standard deviation of log-transformed pfu/ml data from five independent biological replicates.

Although HD100 reduced ML35-irregular populations, we detected some prey population recovery after 72 hours of co-culture, with high variation in concentration (Figure 3). We also observed some reversion to the circular colony morphotype on 48-h and 72-h plates. This contrasts with prey-only controls and co-cultures with NC01, in which the irregular morphotype persisted as the sole colony morphotype over 72 hours. The level of reversion when ML35-irregular was co-cultured with HD100 was highly variable, with proportions of the circular colony morphotype ranging from 0 – 100% across replicates at both 48 and 72 hours (data not shown).

### ML35 lines isolated from NC01 co-cultures carry *rfbG* mutations

We generated short-read sequencing data of ML35-irregular and ML35-circular lines established during initial predation efficiency assays (see above), then used *breseq* to compare reads to the ancestral *E. coli* ML35 genome. This analysis revealed that each line carries a mutation in *rfbG*, annotated as dTDP-rhamnosyl transferase involved in synthesizing the O-antigen component of outer membrane LPS [12]. In two ML35-irregular lines, an IS5-like transposon inserted within *rfbG*, albeit at different nucleotide positions in each line. In the third line, a single nucleotide mutation introduced a premature stop codon four amino acids upstream of the predicted stop. Two ML35-circular lines carry the same single nucleotide mutation resulting in an asparagine to lysine change in amino acid 80. The third line carries a single nucleotide mutation within the predicted stop codon, which altered it to a lysine residue.

## DISCUSSION

Here, we show that predation resistance can emerge rapidly during short-term co-culture of *E. coli* ML35 with *Bdellovibrio* sp. NC01. Prey populations initially decline but subsequently recover despite the continued presence of active predators, and this recovery is correlated with the emergence and eventual predominance of an irregular colony morphotype. ML35-irregular is resistant to killing by NC01 but remains susceptible to *B. bacteriovorus* HD100, indicating that the resistance phenotype is predator-specific rather than a general defense against *Bdellovibrio* predation.

The emergence of the ML35-irregular morphotype observed here extends Varon’s report of a predation-resistant subpopulation arising during co-culture [7]. Whereas Varon observed predation-resistant colony variants after six days of continuous culture of bioluminescent prey with marine predatory bacteria, the predation-resistant *E. coli* morphotype reported here emerged within 24 hours of exposure to *Bdellovibrio* sp. NC01 in batch culture. These studies suggest that predation-resistant subpopulations can arise on relatively short timescales across a range of predator-prey systems and experimental conditions.

Our findings also build on observations by Shemesh and Jurkevitch, who described prey population recovery during short-term co-culture of *Erwinia* prey and *Bdellovibrio* SNE [8]. In their study, surviving *Erwinia* demonstrated resistance to a range of predatory strains, but this resistance phenotype disappeared when predation pressure was removed. By contrast, the ML35-irregular morphotype described here was resistant to killing by NC01 but remained susceptible to HD100, indicating that predation resistance can be specific to particular predator-prey combinations. In addition, ML35-irregular retained both its colony morphotype and predation-resistance phenotype through multiple rounds of streaking for isolation and subsequent growth in liquid culture, all in the absence of predation pressure. These studies show that predation resistance that develops during short-term co-culture can vary in both persistence and predator specificity.

Whole-genome sequencing of clonal lines of both *E. coli* colony morphotypes established from three independent co-cultures revealed that each line carries a mutation in *rfbG*, which is predicted to encode a GT-A fold glycosyltransferase involved in O-antigen synthesis. Two ML35-irregular lines carry IS element insertions within the *rfbG* coding sequence, which likely disrupt the gene, and the third carries a premature stop codon four residues from the C-terminus. In other GT-A fold enzymes, C-terminal residues are required for catalytic activity [13], suggesting this mutation may impair function. Two ML35-circular lines carry the same missense mutation (N80K), and the third carries a nonstop mutation. The effects of these mutations on RfbG function are unknown. Overall, these data identify *rfbG* as a recurrent mutation target among *E. coli* ML35 recovered from NC01 co-cultures, although additional experiments are needed to determine whether these mutations contribute to predation resistance and whether NC01 predation is the selective force.

Given the predicted function of RfbG, our findings also raise the possibility that O-antigen contributes to *E. coli* ML35 susceptibility to predation by NC01. Under this hypothesis, disruption of *rfbG* affects O-antigen synthesis, leading to loss of O-antigen from the outer membrane and resistance to predation by NC01. If supported, this mechanism would differ from prior work showing protective effects of O-antigen against *Bdellovibrio* predation. *Vibrio cholerae* mutants lacking O-antigen were attacked more rapidly by *B. bacteriovorus* [14], and restoration of O-antigen synthesis in *E. coli* K-12 reduced susceptibility to some predatory strains [6]. The relationship between O-antigen and susceptibility to predation may depend on the specific predator-prey combination. Further investigation, including phenotypic testing of ML35-circular isolates, genetic analysis of *rfbG*, and characterization of LPS structure, will clarify the mechanisms underlying the predation resistance phenotype described here.

## ACKNOWLEDGMENTS

Research reported in this publication was made possible by the use of services available through the Rhode Island INBRE Centralized Research Core Facility (RRID:SCR_028658) and supported in part by the Rhode Island Institutional Development Award (IDeA) Network of Biomedical Research Excellence from the National Institute of General Medical Sciences of the National Institutes of Health under grant number P20GM103430.

